# Atmospheric nitrogen fixation by eukaryotes: Should we reconsider the nitrogen cycle in nature?

**DOI:** 10.1101/2022.09.29.510055

**Authors:** Nikolaos Arapitsas, Christos A. Christakis, George Angelakis, Aristeidis Maragkoudakis, Kiriakos Kotzabasis, Panagiotis F. Sarris

## Abstract

Nitrogen is the most abundant element in our planet’s atmosphere. Approximately 78% of the atmosphere is made up of nitrogen gas (N2). It is essential for organisms, as a major component of biomolecules and cellular biochemistry. Until today, atmospheric N_2_ assimilation in eukaryotic organisms has been reported to occur only through specific symbiotic N_2_-fixing bacterial groups that make it available to plants, the base of the food chain.

In this work, using Gas Chromatography with Thermal Conductivity Detection, we proved the direct consumption of atmospheric N_2_ by eukaryotic organisms. Three yeast species: *Debaryomyces hansenii, Metschnikowia reukaufii* and model-species *Saccharomyces cerevisiae* are able to directly assimilate atmospheric nitrogen in various conditions, with a yet unknown mechanisms that are not based on the presence of known N_2_-fixation enzymes. Our findings could change the way we consider the N_2_-assimilation in nature, laying the basis for future focused studies on the molecular mechanism underling the biological phenomenon of eukaryotic N_2_-assimilation.

## Introduction

Nitrogen is the most abundant element in Earth’s atmosphere, with its molecular form (dinitrogen or N_2_) making up of approximately 78% of the total mass. It is a necessary element for all living organisms as an essential component of proteins, nucleic acids and in the energy transfer molecule of ATP (adenosine triphosphate). Despite its high concentration in the atmosphere, N_2_ is biochemically unavailable to most organisms and especially the eukaryotes. For example plants and animals do not express the required enzymes (nitrogenases) to make use of atmospheric nitrogen. For this, nitrogen has to be converted to a more usable forms. There are two ways that nitrogen can be fixed to be useful for living organisms: 1. <u>B</u>iological <u>N</u>itrogen <u>F</u>ixation (BNF). In this way the nitrogen gas enters the soil where it is absorbed by specific members of several bacterial and archaeal phyla, which convert it to ammonium ions (NH_4_^+^). Ammonium ions can be used by plants and this is the way that the vast majority of the available nitrogen to organisms, enters the food web and becomes part of the N_2_ cycle; 2. A small amount of nitrogen can also enter the food web through lightning, which converts atmospheric nitrogen into ammonia and nitrate (NO_3_) that enter soil with rainfall.

The first report of BNF, in 1888 by Hellriegel and Willfarth [1,2], was followed by series of investigations related to atmospheric N_2_ fixation by living organisms which reported that this unique characteristic of biological N_2_ assimilation is limited to specialised groups of soil and endophytic bacteria[3–6]. While, the conversion of atmospheric N_2_ into bioavailable ammonia by the enzyme nitrogenase, was reported 1.2 centuries after the first description of BNF [4,7].

However, for an interval of more than a century, several research groups have tried to uncover evidence of atmospheric nitrogen fixation by a number of eukaryotes [3]. The experimental designs included the use of various model organisms, and the detection of the N_2_-fixation process indirectly, via the measurement of the nitrogen containing by-products of the process. The results of these studies were both positive and negative [3]. Despite the research of atmospheric N_2_ assimilation by eukaryotes no definite proof of the process has been put forward as yet.

Yeasts are unicellular eukaryotic microorganisms classified as members of the kingdom Fungi and belong to the subkingdom *Dikarya*. At least 1,500 species are recognized up today, including the well-studied model-organism *Saccharomyces cerevisiae* [8–11]. Select yeast species can colonize the inner compartments of plants and spend part of their lifecycle as endophytes without causing damage to their hosts [11,12]. Endophytic yeasts have been reported to be a viable way of reducing fertilizer inputs in agriculture or as biocontrol agents against plant pathogens [13–15]. However, many aspects of endophytic yeasts lifecycle still need to be elucidated, especially on their means of plant host colonization and their production of viable components [11].

In this study, we identified and examined the ability of three yeast species to assimilate the atmospheric N_2_ using for the first time an experimental approach which enabled us to observe the direct atmospheric N_2_ assimilation.

## Results and Discussion

### Identification of yeast species that grow in N2-free culture media

We screened more than 600 endophytic microorganisms, obtained from <u>c</u>rop <u>w</u>ild <u>r</u>elative (CWR) halophytes for their N_2_-fixation ability, through cultivation on the solid nitrogen-free media NFb [16] and <u>N</u>itrogen-<u>f</u>ree - <u>M</u>inimal <u>S</u>alt <u>M</u>edium with amendments (MSM-NF), which are widely used to screen for nitrogen-fixing microbes. Apart from several bacterial species that we found to grow well on these media, we identified two endophytic yeast species that grew well on NFb and MSM-NF (Fig. 1 and Suppl. Fig 1). Based on their ITS1/2 sequence they have been characterised as *Debaryomyces hansenii* and *Metschnikowia reukaufii. Debaryomyces hansenii* is one of the most representative endophytic yeast species [13,17], while *Metschnikowia reukaufii* is a floricolous *Saccharomycetales* yeast, which as an endophyte is usually found dwelling in nectar and fruit [18].

**Figure 1.**
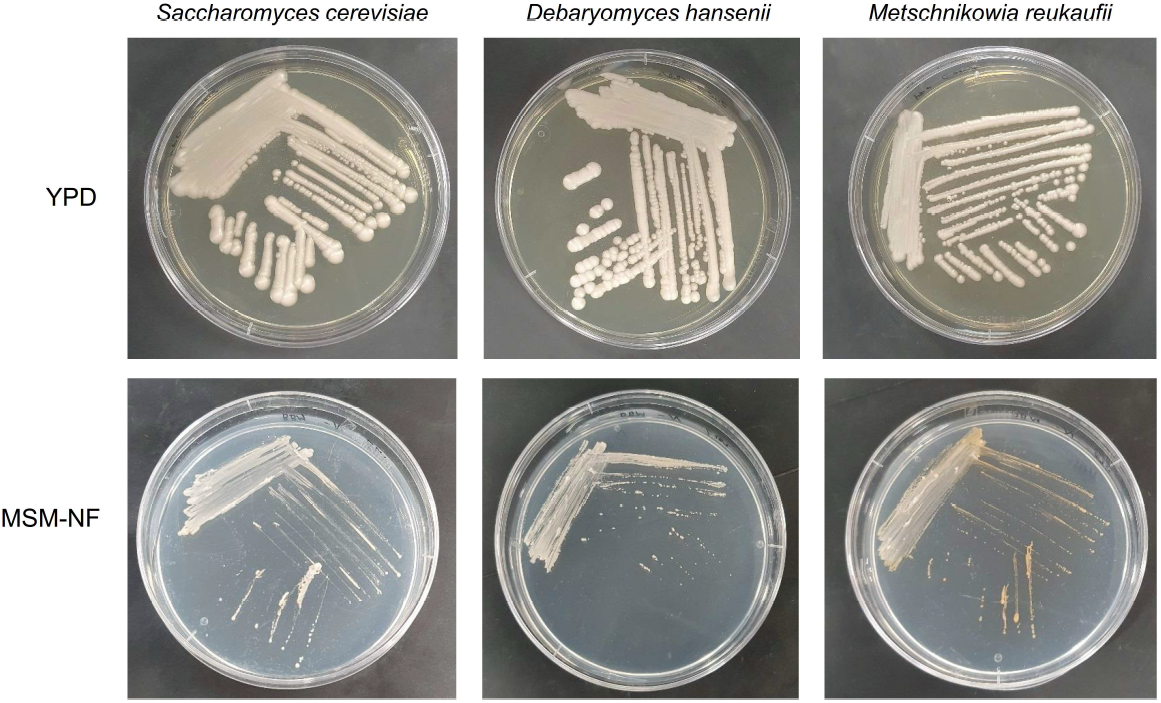
Growth of *Saccharomyces cerevisiae, Debaryomyces hansenii* and *Metschnikowia reukaufii* on rich medium (YPD; 6dpi) and on <u>N</u>itrogen-<u>f</u>ree - <u>M</u>inimal <u>S</u>alt <u>M</u>edium with amendments (MSM-NF; 6dpi).

Based on their growth on nitrogen starved media, we hypothesized that the yeast isolates have the ability to consume atmospheric N_2_, since no other nitrogen source was available. To *in-depth* investigate our initial observation, we continued cultivating the endophytic yeasts on solid NFb for multiple generations to control for potential intracellular stored nitrogen compounds.

### Yeast species do not express known N_2_-fixation enzymes

To control for the potential presence of the nitrogenase enzymes that are responsible for prokaryotic nitrogen fixation in yeast species, we cultivated the yeasts on NFb plates that lacked molybdenum (Mo) sources, on plates that lacked iron (Fe) sources and on plates lacking both Mo and Fe (Suppl. Fig. 7) [19]. As it has been extensively reported, the Mo-nitrogenases are responsible for most BNF in nature [19,20]. Furthermore, we performed cultivation on semi-solid NFb media [21] and on solid MSM-NF medium (Fig.1 and Suppl. Fig. 1). We also successfully cultivated all yeast species on solid <u>n</u>itrogen-<u>f</u>ree <u>m</u>ineral broth <u>m</u>edium (NFMM), based on a 2013 report on successful cultivation of yeasts NFMM medium [15].

Furthermore, genome mining studies did not reveal the presence of nitrogenase-like enzymes in the genome of these yeast species, a finding that explains their ability to grow, even slowly, on minimal media lacking Mo and Fe (Suppl. Fig. 6). These findings also are in consistent with previous studies reporting the absence of nitrogenase-like activity in *Saccharomyces* species [22].

### *Using* GC-TCD *we directly observed that yeasts consume atmospheric N_2_*

To *in-depth* investigate our initial observations and after all the successful cultivation experiments in solid N_2_-free media, we proceeded in liquid cultivation experiments in order to track the growth of the yeast and the consumption of N_2_ and O_2_ gases. The gas consumption was tracked with the <u>G</u>as <u>C</u>hromatography with a <u>T</u>hermal <u>C</u>onductivity <u>D</u>etector (GC-TCD) analysis method. Apart from the two endophytic yeasts we also employed the model-species *Saccharomyces cerevisiae[23]* to investigate if the atmospheric N_2_-assimilation is generalized among the *Saccharomycetales* order. Furthermore, we included as biological negative controls the bacteria *Escherichia coli* and *Rhizobium laguerreae. E.coli* strains lack nitrogenase genes and thus incapable of nitrogen fixation. Whereas *R. laguerreae* is a well-studied N_2_-fixing species, which however has an oxygen sensitive nitrogenase and the N_2_-fixation ability is inducible only upon host-plant colonization and formation of the specialised root-nodules in various legume species [24,25].

All organisms were cultivated in biological reactors, i.e. in 120 mL serum bottles containing 50 mL liquid media capped with butyl rubber stoppers and with a fixed headspace (70 mL of atmospheric air). Technical blank controls of all assays were included that contained uninoculated media. Liquid cultivations were performed using both nitrogen-free minimal media (mannitol NFMM, glucose NFMM, NFb), as well as in rich media (YPD) [26].

The GC-TCD assays in the reactors containing yeasts, revealed, for the first time in the literature, the direct consumption of atmospheric N_2_ by eukaryotic organisms (Fig. 2). The inclusion of blank controls clearly indicated that the observed atmospheric N_2_-assimilation is a biological phenomenon (Fig. 2 and Suppl. Fig. 2). Interestingly, all the examined yeast species, including the model-species *Saccharomyces cerevisiae*, were able to consume the atmospheric N_2_ (Fig. 2). The different examined species revealed differential speed of atmospheric N_2_-assimilation (Fig. 2). These findings are consistent with previous studies reporting that various yeast species are able to produce NH_4_^+^ in the absence of a nitrogen source in the culture media [15].

**Figure 2.**
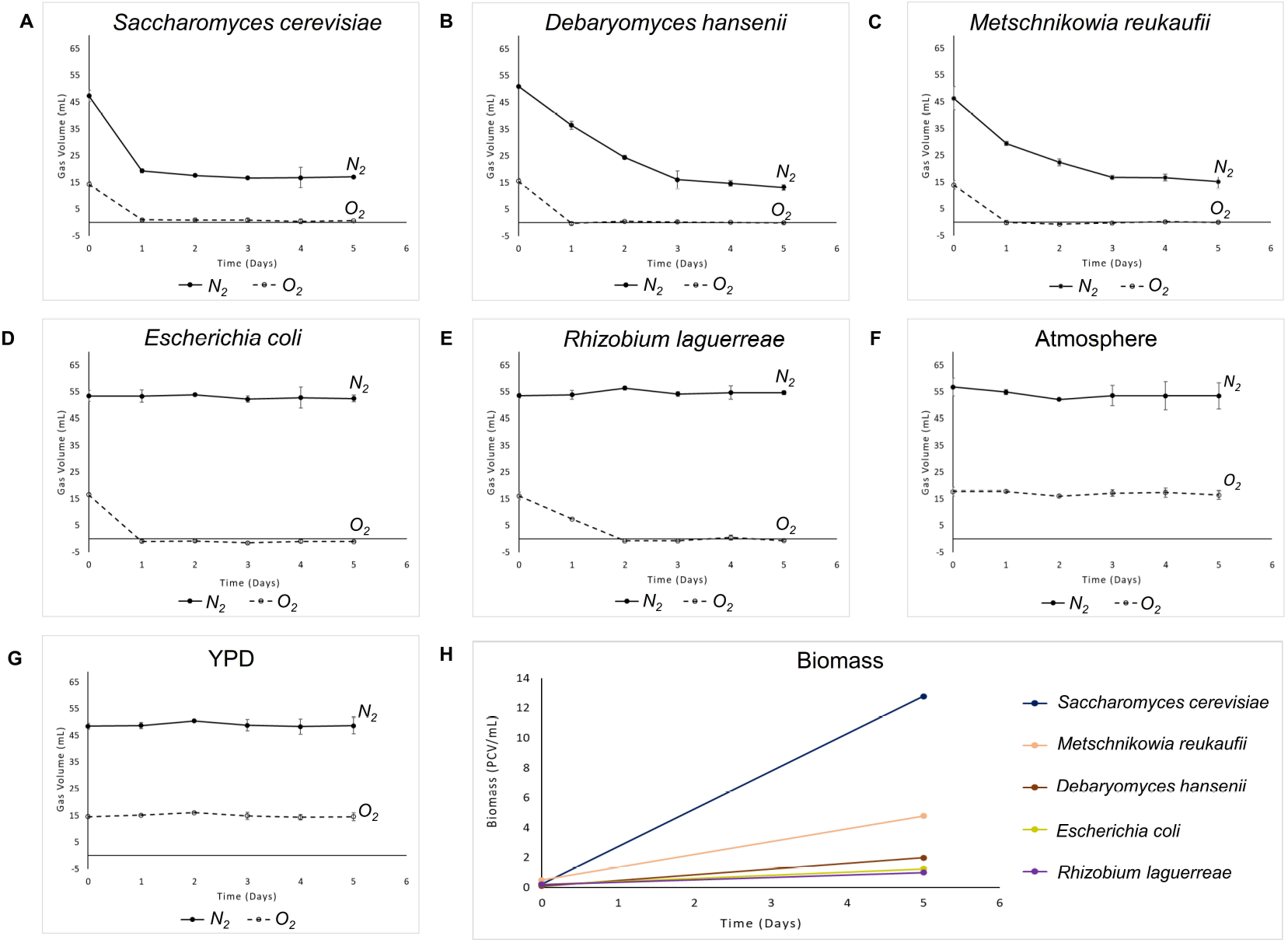
GC-TCD measurements of N_2_-assimilation (solid line) and O_2_ (dashed line) in closed liquid cultures in YPD medium: **A.** of *Saccharomyces cerevisiae*; **B.** of *Debaryomyces hansenii*; **C.** of *Metschnikowia reukaufii*; **D.** negative control *E. coli*; **E.** negative control *Rhizobium laguerreae*; **F.** negative control Atmosphere; **G.** N_2_-assimilation (solid line) and O_2_ levels (dashed line) in un-inoculated YPD. **H.** Biomass growth graph of all species in this assay.

### Yeasts are able to consume atmospheric N_2_ in presence or absence of O_2_ and independently of a nitrogen source in the medium

Cultivation experiments in rich media and with differing atmospheric conditions revealed that the examined yeast species are able to utilize the atmospheric N_2_ regardless of the presence of a nitrogen source in the medium (media amended with NH_4_^+^ or NO_3_^-^) (Suppl. Fig. 3 & 4). Furthermore, all yeast species demonstrated unabated ability to atmospheric N_2_-consumption, in either presence or absence of O_2_ (Fig. 3 & Suppl. Fig. 5).

**Figure 3.**
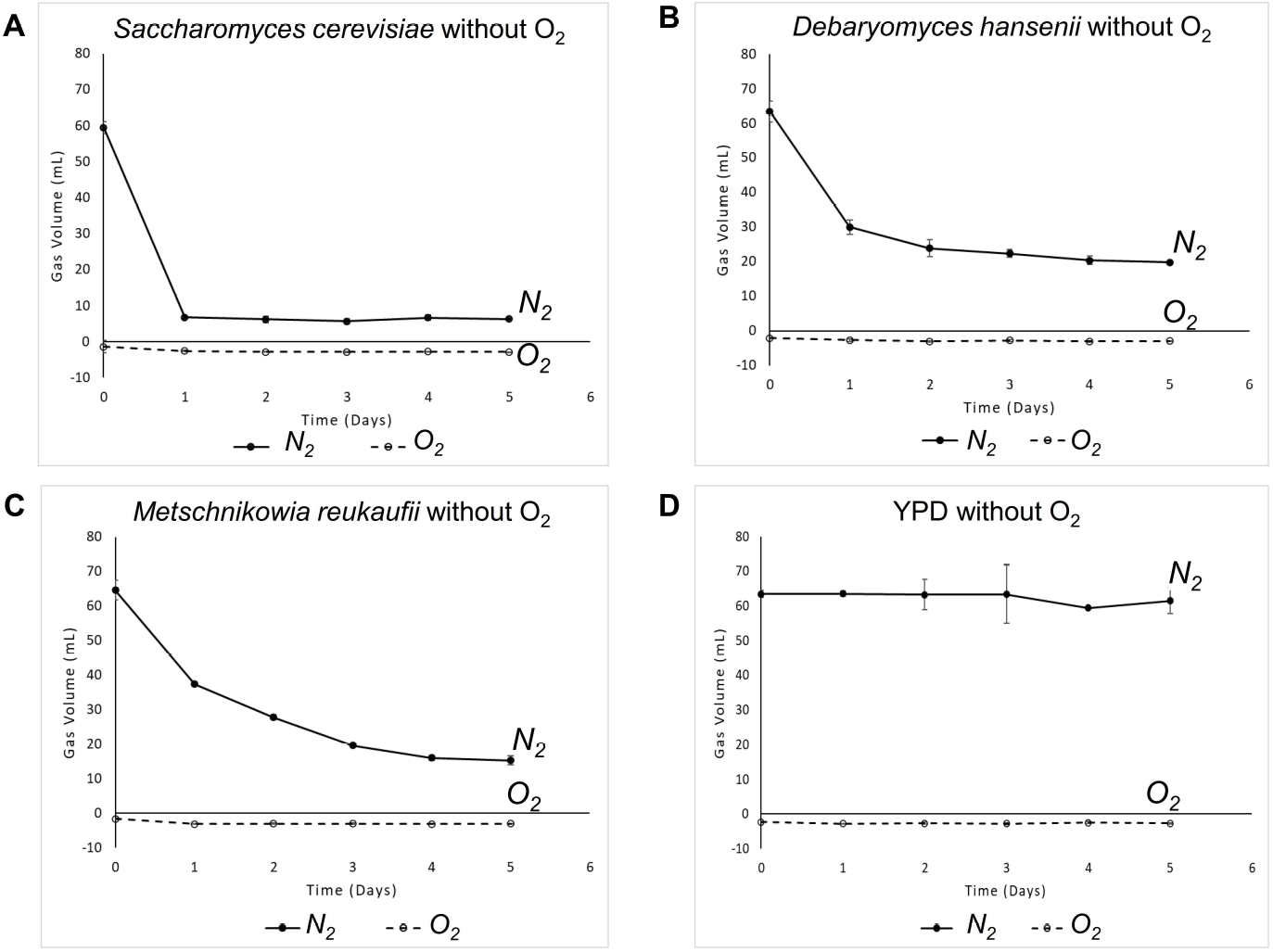
GC-TCD measurements of N_2_-assimilation (solid line) and O_2_ (dashed line) in yeast closed liquid cultures in YPD medium without O_2_ on the headspace. **A.** *Saccharomyces cerevisiae* in YPD medium without O_2_; **B.** *Debaryomyces hansenii* in YPD medium without O_2_; **C.** *Metschnikowia reukaufii* in YPD medium without O_2_; **D.** Negative control: YPD medium without O_2_.

To rule out any potential presence of prokaryotic endosymbionts that are able to consume N_2_[22,27], we cultivated all the examined yeast strains in a cocktail of antibiotics (Suppl. Fig. 7). This treatment did not affect the ability of the examined yeast species to atmospheric N_2_-assimilation (Suppl. Figs 8 & 9). Furthermore, we performed 16S and ITS diagnostic PCRs in order to rule out any possibility of endosymbionts (Suppl. Fig. 10).

### Laboratory yeast strains are also able to consume atmospheric N_2_

In order to investigate if this biological phenomenon has been maintained in laboratory yeast strains, different haploid *S. cerevisiae* lab strains were tested for their ability to atmospheric N_2_-assimilation. All the examined strains revealed positive results (Suppl. Figs 11 & 12).

## Conclusion

The available supply of nitrogen is a limitation in natural ecosystems which controls many processes, such as primary production and decomposition. This indicates that nitrogen is often the limiting nutrient, which in short supply limits the growth of organisms or populations. However, having nitrogen around is different from being able to make use of. The eukaryotic organisms, including plants and animals, have no biochemical mechanisms to convert atmospheric N_2_ into an assimilable form, since, they don’t have the right enzymes to capture, or fix, atmospheric nitrogen. So, up today it is proposed that the only way N_2_ to enter the ecosystems as a part of the food web of chemicals, was the atmospheric N_2_-fixation by prokaryotes. Some of these N_2_-fixing bacterial species are free-living in soil or water, while others are beneficial plant symbionts.

In this work, our data clearly indicate the presence of a novel, yet unknown, mechanism of atmospheric N2-assimilation in yeast. Interestingly, some of the examined yeast species appeared to be cosmopolitan endophytic in plants. This finding is groundbreaking, since, it could change the way we consider the N2 cycle in nature so far. Furthermore, the potential biotechnological applications in food production and elsewhere, seem to be enormous. We strongly believe that our findings establish the basis for future focused studies on the identification of the eukaryotic N2-assimilation mechanism.

## Materials and Methods

### Isolation of Debaryomyces hansenii and Metschnikowia reukaufii from plant material

*Debaryomyces hansenii* and *Metschnikowia reukaufii* were isolated from surface-sterilized *Matthiola tricuspidata* leaves and *Teucrium capitatum* roots, respectively, as it was described previously [28]. After sterilization, 100 μL of solution from the last wash was plated as a negative control. The extract obtained from the surface-sterilized plant material was plated. The plates were incubated for 4 - 7 d at 28°C and single colonies of the yeast species were selected and repeatedly streaked onto NA plates (5 g/L bacto-peptone, 3 g/L yeast extract, 5 g/L NaCl, 15 g/L agar, pH 7.4) to obtain pure cultures.

### Identification of yeast isolates

To identify the yeast isolates, the Internal Transcribed Spacer Regions 1 and 2 (ITS1/2) were sequenced using the Sanger sequencing method. The crude genomic DNA extraction from yeasts was performed using the Chelex® 100 sodium form (50-100 mesh) (Sigma-Aldrich, Darmstadt, Germany). Colonies from each yeast culture grown on YPD agar medium (10 g/L yeast extract, 20 g/L bacto-peptone, 20 g/L glucose, 20 g/L agar, pH 6.0-6.5) was diluted in 40 μL ethanol 50% v/v. 200 μL of an aquatic suspension of Chelex® 100 10% w/v was added and followed by brief vortex. The samples were heated at 98°C for 20 min and centrifuged at 18,000 × g for 10 min. A volume equal to 2 μL of the supernatant was used as a template in a PCR reaction using primers ITS1F: 5’-CTTGGTCATTTAGAGGAAGTAA-3’ and ITS4: 5’-TCCTCCGCTTATTGATATGC-3’ to amplify the ITS1/2 sequence [29]. The template was mixed with 1× Taq Pol Buffer (Minotech, Heraklion, Greece), 1.5 mM MgCl_2_, 0.2 mM dNTPs, 0.5 μM of ITS1F and ITS4 primers, 0.04 U Taq DNA Pol (Minotech, Heraklion, Greece), and ultrapure H_2_O up to a total volume of 25 μL. PCR reactions were amplified in a BioRad T-100 Thermocycler (Bio-Rad Laboratories) with initial denaturation at 94°C for 2 min, followed by 30 cycles of 45 sec at 94°C, 30 sec annealing at 55°C, 90 sec extension at 72°C, and a final extension step at 72°C for 10 min. PCR products were purified using the NucleoSpin® Gel and PCR Clean-up kit (Macherey-Nagel™, Düren, Germany) according to the manufacturer’s instructions. The DNA was eluted twice in 25 μL ultrapure H_2_O. The purified PCR products were sequenced by Genewiz (Germany, Leipzig) with the ITS1F and ITS4 primers. The resulting sequences of the ITS1/2 region were inspected using the SnapGene® software (from Insightful Science; available at snapgene.com) and the start/end regions of low quality were manually trimmed o□. The trimmed ITS1/2 sequences were queried in BLAST using the NCBI GenBank database for identification and documentation of the described yeast isolate with the closest sequence similarity.

### Yeast Growth in Nitrogen-Free Media

To investigate growth on nitrogen-free media, *Debaryomyces hansenii, Metschnikowia reukaufii* and *Saccharomyces cerevisiae* were streaked onto three distinct Glucose Nitrogen-Free Media: NFb (5 g/L malic acid, 0.5 g/L K_2_HPO_4_, 0.2 g/L MgSO_4_·7H_2_O, 0.1 g/L NaCl, CaCl_2_·2H_2_O 0.02 g/L, 2 mL micronutrient solution (0.4 g/L CuSO_4_·5H_2_O, 0.12 g/L ZnSO_4_·7H_2_O, 1.4 g/L H_3_BO_3_, 1.0 g/L Na_2_MoO_4_·2H_2_O, 1.5 g/L MnSO_4_·H_2_O), 2 mL Bromothymol blue solution (0.5% w/v in 0.2 N KOH), 4 mL Fe(III)-EDTA (1.64% w/v), 1 mL vitamin solution (100 mg/L biotin, 200 mg/L pyridoxal-HCl), 15 g/L agar, pH 6.8), Nitrogen-free Minimal Salt Medium with amendments (MSM-NF) (25 mg/L CaCl_2_·2H_2_O, 75 mg/L MgSO_4_·7H_2_O, 75 mg/L K_2_HPO_4_, 175 mg/L KH_2_PO_4_, 25 mg/L NaCl, 11.42 mg/L H_3_BO_3_, 0.5 mL EDTA solution (100 g/L EDTA, 62 g/L KOH), 0.5 mL acidified iron solution (9.96 g/L FeSO_4_·7H_2_O, 0.192% v/v H_2_SO_4_), 0.5 mL trace metal solution (17.64 g/L ZnSO_4_·7H_2_O, 2.88 g/L MnCl_2_·4H_2_O, 1.42 g/L MoO_3_, 3.14 g/L CuSO_4_·5H_2_O), 15 g/L agar, pH 6.0-6.2), and NFMM (1 g/L K_2_HPO_4_, 1 g/L CaCl_2_, 0.5 g/L NaCl, 0.25 g/L MgSO_4_·7H_2_O, 0.01 g/L FeSO_4_·7H_2_O, 0.01 g/L Na_2_MoO_4_·2H_2_O, 0.01 g/L MnSO_4_·5H_2_O, 7 g/L glucose, 20 g/L agar), and incubated at 28°C for 5 d. For the NFb medium without Mo and Fe, Na_2_MoO_4_ and Fe(III)-EDTA were omitted, respectively.

### Preparation of liquid cultures of yeasts and bacterial species for GC-TCD measurements

*Debaryomyces* hansenii, *Metschnikowia reukaufii* and *Saccharomyces cerevisiae* were initially grown on YPD agar medium and incubated at 28°C for 3 – 5 d. For the laboratory strains of *Saccharomyces cerevisiae* AH109 and PJ69-4A, the YPD medium was supplemented with 30 mg/L adenine (YPDA). The bacteria *Escherichia coli* DH10B and *Rhizobium laguerreae* were grown on LB agar (10 g/L tryptone, 5 g/L yeast extract, 10 g/L NaCl, 15 g/L agar) and NA medium, respectively. For the liquid cultures, colonies from the plates were used to inoculate the appropriate medium in liquid form (YPD, LB or NA without agar). The cell concentration was measured by centrifugation at 3,000 × g for 5 min in <u>p</u>acked <u>c</u>ell <u>v</u>olume (PCV) tubes. The culture volume was adjusted to 50 mL with a cell concentration of 0.2 – 0.5 PCV/mL. Every culture was transferred in 120 mL serum bottles capped with butyl rubber stoppers and with a fixed headspace of 70 mL, and incubated at 28°C, 200 rpm for 5 – 8 d. Technical blank controls were included that contained uninoculated media. Both the serum bottles and the rubber stoppers were treated with 1 M HCl for 3 h and washed thoroughly with ultrapure H_2_O prior to sterilization to eliminate any mineral nitrogen residue in the vials. The biomass of the cultures expressed in PCV/mL was measured at the end of their incubation period by the centrifugation of a sample from each culture in PCV tubes at 3,000 × g for 5 min.

### Cultivation experiments in media supplemented with nitrogen sources and without O_2_

Liquid cultures of *Debaryomyces* hansenii, *Metschnikowia reukaufii* and *Saccharomyces cerevisiae* in YPD medium supplemented with NH_4_^+^ (YPD with 0.25% w/v (NH_4_)_2_SO_4_) and NO_3_^-^ (YPD with 0.5% w/v NaNO_3_) were prepared to evaluate the effect of nitrogen availability in the nitrogen-assimilation ability of the yeasts. Additionally, to generate an atmosphere without oxygen in the headspace of the cultures, two syringes were used to fix the headspace atmosphere. One of the syringes supplied the headspace with nitrogen gas from a tank, while, simultaneously, the other syringe removed the pre-existing atmosphere headspace for 30 s. By this method, a headspace without oxygen-filled with nitrogen gas was created.

### GC-TCD measurements of N_2_ and O_2_ in closed liquid cultures

Oxygen and nitrogen measurements were made by <u>g</u>as <u>c</u>hromatography, using a <u>t</u>hermal <u>c</u>onductivity <u>d</u>etector (GC-TCD) (Shimadzu GC 2010 Plus, Kyoto, Japan). To separate O_2_ and N_2_, argon was used as the carrier gas under pressure of 5 bars and at an oven temperature of 120 °C. A capillary Vici Metronics MC (Poulsbo, USA) column was used, with length 30 m (diameter: 0.53 mm) and film thickness 20 μm. The temperature of TCD was set at 200 °C for the detector and at 180 °C for the injector. Samples of volume equal to 200 μL were collected from the headspace using a gas-tight syringe and injected into the GC-TCD to obtain the measurements. The headspace sampling from each culture was repeated at least three times, unless specified. Atmospheric samples of equal volume and samples from the headspace of uninoculated media served as blank controls. For each culture, O_2_ and N_2_ GC-TCD measurements of the headspace were taken every day, unless specified. The quantification of N_2_ and O_2_ volume was done by injecting known quantities in the GC-TCD. The mean value and the standard deviation of N_2_ and O_2_ volumes in the headspace of the closed bottles, were calculated for each time point and for every serum bottle, based on the GC-TCD measurements, and used to plot the change in the content of N_2_ and O_2_ in the culture headspace over time.

### Antibiotic media to rule out the presence of prokaryotic endosymbionts in yeast species

All the yeast species were cured for potential endobacteria as described in Sen et al. (2019) [30]. In short, yeast cells were inoculated three times in YPD medium in the presence of tetracycline (10 μg/mL), gentamycin (30 μg/mL), and ampicillin (100 μg/mL) to obtain prokaryote-free yeast strains. Single colonies of all yeast species were inoculated again on YPD plates without antibiotics. The obtained strains were used in liquid cultures in closed atmosphere using GC-TCD measurements of N_2_ and O_2_ as described previously. No effect was observed in their ability to consume the atmospheric N_2_.

### Diagnostic PCR for ITS1/2 and 16S rDNA sequences

The crude DNA of the yeast species was used as template for diagnostic PCR reactions specific for the ITS1/2 and the 16S rDNA to rule out the presence of prokaryotes in the yeast cultures. The primers ITS1F and ITS4 were used for the ITS1/2 region, and the primers 27F: 5’-AGAGTTTGATCCTGGCTCAG-3’ and 1492R: 5’-GTTTACCTTGTTACGACTT-3’ were used for the 16S rDNA [31]. Apart from the three yeast species, *E*. *coli* DNA (positive control for 16S rDNA), fungal DNA (positive control for ITS1/2 region) and negative control (reaction without template) were also used as templates for both reactions. The PCR amplification was prepared as described previously. For the electrophoresis of the PCR products, 5 μL of each reaction were mixed with 1 μL 6× Orange G, loaded on an 1.2% w/v agarose gel and ran at 90 V for 30 min. The DNA was visualized under UV transillumination.

## Supplementary Figure Legends

**Suppl. Figure 1**. Growth of *Saccharomyces cerevisiae, Debaryomyces hansenii* and *Metschnikowia reukaufii* on nitrogen-free NFb medium (carbon source: malate; 4dpi).

**Suppl. Figure 2**. GC-TCD chromatograms of N_2_ and O_2_ in closed liquid cultures in YPD medium at Day 0 and Day 5 after the inoculation: **A.** of Saccharomyces cerevisiae; **B.** of *Debaryomyces hansenii*; **C.** of *Metschnikowia reukaufii*; **D.** negative control *E. coli*; **E.** negative control *Rhizobium laguerreae*; **F.** negative control Atmosphere. Three independent replicates are presented in each panel. Y axis: Intensity.

**Suppl. Figure 3**. GC-TCD measurements of N_2_-assimilation (solid line) and O_2_ (dashed line) in yeast closed liquid cultures in YPD medium supplemented with NH_4_^+^ or NO_3_^-^: **A.** *Saccharomyces cerevisiae* supplemented with NH_4_^+^; **B.** *Debaryomyces hansenii* supplemented with NH_4_^+^; **C.** *Metschnikowia reukaufii* supplemented with NH_4_^+^; **D.** *Saccharomyces cerevisiae* supplemented with NO_3_^-^; **E.** *Debaryomyces hansenii* supplemented with NO_3_^-^; **F.** *Metschnikowia reukaufii* supplemented with NO_3_^-^.

**Suppl. Figure 4**. GC-TCD chromatograms of N_2_ and O_2_ in yeast closed liquid cultures in YPD medium supplemented with NH_4_^+^ or NO_3_^-^ at Day 0 and Day 5 after the inoculation: **A.** *Saccharomyces cerevisiae* supplemented with NH_4_^+^; **B.** *Debaryomyces hansenii* supplemented with NH_4_^+^; **C.** *Metschnikowia reukaufii* supplemented with NH_4_^+^; **D.** *Saccharomyces cerevisiae* supplemented with NO_3_^-^; **E.** *Debaryomyces hansenii* supplemented with NO_3_^-^; **F.** *Metschnikowia reukaufii* supplemented with NO_3_^-^. Three independent replicates are presented in each panel. Y axis: Intensity.

**Suppl. Figure 5**. GC-TCD chromatograms of N_2_ and O_2_ in yeast closed liquid cultures in YPD medium without O_2_ on the headspace at Day 0 and Day 5 after the inoculation: **A.** *Saccharomyces cerevisiae* in YPD medium without O_2_; **B.** *Debaryomyces hansenii* in YPD medium without O_2_; **C.** *Metschnikowia reukaufii* in YPD medium without O_2_; **D.** Negative control: YPD medium without O_2_.

Three independent replicates are presented in each panel (two independent replicates are presented for *Saccharomyces cerevisiae* in YPD medium without O_2_ at Day 0). Y axis: Intensity.

**Suppl. Figure 6**. Growth of all yeast species on nitrogen free NFb medium lacking molybdenum (Mo) sources (NFb w/o Mo), iron (Fe) sources (NFb w/o Fe) and both Mo and Fe sources (NFb w/o Mo and Fe); 4dpi.

**Suppl. Figure 7**. Growth of all yeast species on reach medium (YPD) supplemented with an antibiotic cocktail; 6dpi.

**Suppl. Figure 8**. GC-TCD measurements of N_2_-assimilation (solid line) and O_2_ (dashed line) in closed liquid cultures of all yeast species upon treatment with antibiotics: **A.** *Saccharomyces cerevisiae*; **B.** *Debaryomyces hansenii*; **C.** *Metschnikowia reukaufii*.

**Suppl. Figure 9**. GC-TCD chromatograms of N_2_ and O_2_ in yeast in closed liquid cultures of all yeast species upon treatment with antibiotics at Day 0 and Day 5 after the inoculation: **A.** *Saccharomyces cerevisiae*; **B.** *Debaryomyces hansenii*; **C.** *Metschnikowia reukaufii*. Three independent replicates are presented in each panel. Y axis: Intensity.

**Suppl. Figure 10**. Electrophoresis of ITS1/2 and 16S diagnostic PCR reactions. ITS1/2: 1. *Saccharomyces cerevisiae, 2. Debaryomyces hansenii*, 3. *Metschnikowia reukaufii*, 4. *Escherichia coli* (negative control for the ITS1/2 reaction), 5. Fungal DNA (positive control for the ITS1/2 reaction), 6. PCR negative control; 16S: 1. *Saccharomyces cerevisiae, 2. Debaryomyces hansenii*, 3. *Metschnikowia reukaufii*, 4. *Escherichia coli* (positive control for the 16S reaction), 5. Fungal DNA (negative control for the 16S reaction), 6. PCR negative control.

**Suppl. Figure 11**. GC-TCD measurements of N_2_-assimilation (solid line) and O_2_ (dashed line) in closed liquid cultures of laboratory yeast strains with and without O_2_ on the headspace: **A.** *Saccharomyces cerevisiae* AH109; **B.** *Saccharomyces cerevisiae* PJ69-4A; **C.** *Saccharomyces cerevisiae* AH109 without O_2_; **D.** *Saccharomyces cerevisiae* PJ69-4A without O_2_.

**Suppl. Figure 12**. GC-TCD chromatograms of N_2_ and O_2_ in yeast in closed liquid cultures of laboratory yeast strains with and without O_2_: **A.** *Saccharomyces cerevisiae* AH109 (at Day 0 and Day 8); **B.** *Saccharomyces cerevisiae* PJ69-4A (at Day 0 and Day 5); **C.** *Saccharomyces cerevisiae* AH109 without O_2_ (at Day 0 and Day 8); **D.** *Saccharomyces cerevisiae* PJ69-4A without O_2_ (at Day 0 and Day 8). Three independent replicates are presented in each panel. Y axis: Intensity.

