## Supplementary Figures for "Atmospheric nitrogen fixation by eukaryotes: Should we reconsider the nitrogen cycle in nature?"

*Saccharomyces cerevisiae* *Debaryomyces hansenii* *Metschnikowia reukaufii*

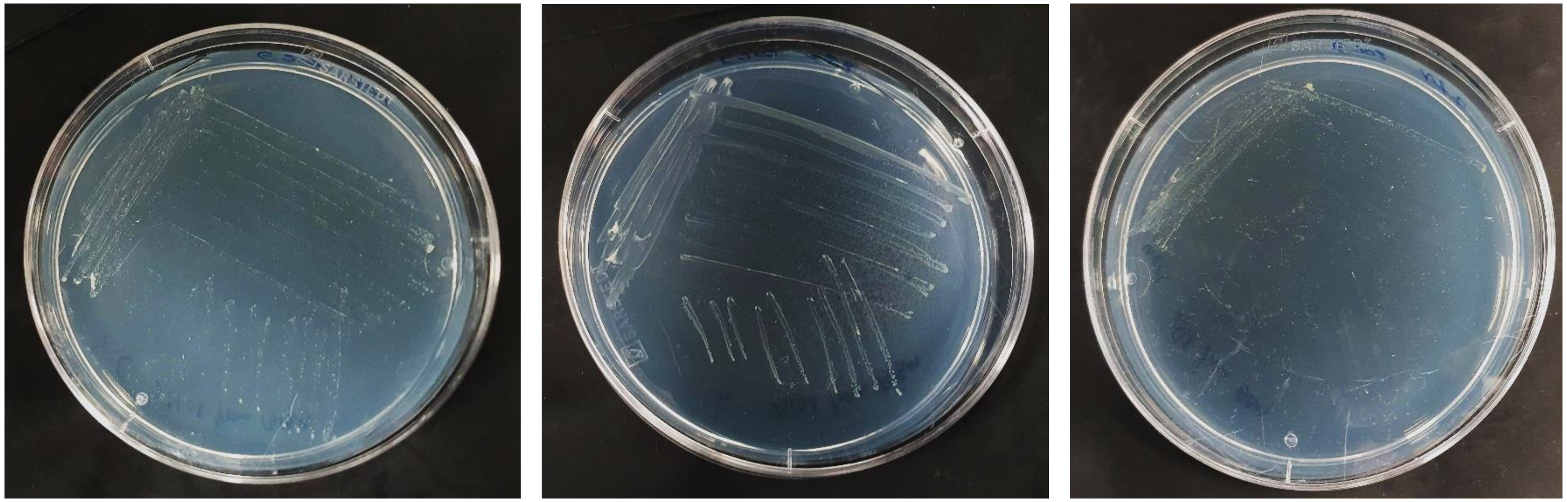

**Suppl. Figure 1.** Growth of *Saccharomyces cerevisiae*, *Debaryomyces hansenii* and *Metschnikowia reuakufii* on nitrogen-free NFb medium (carbon source: malate; 4dpi).

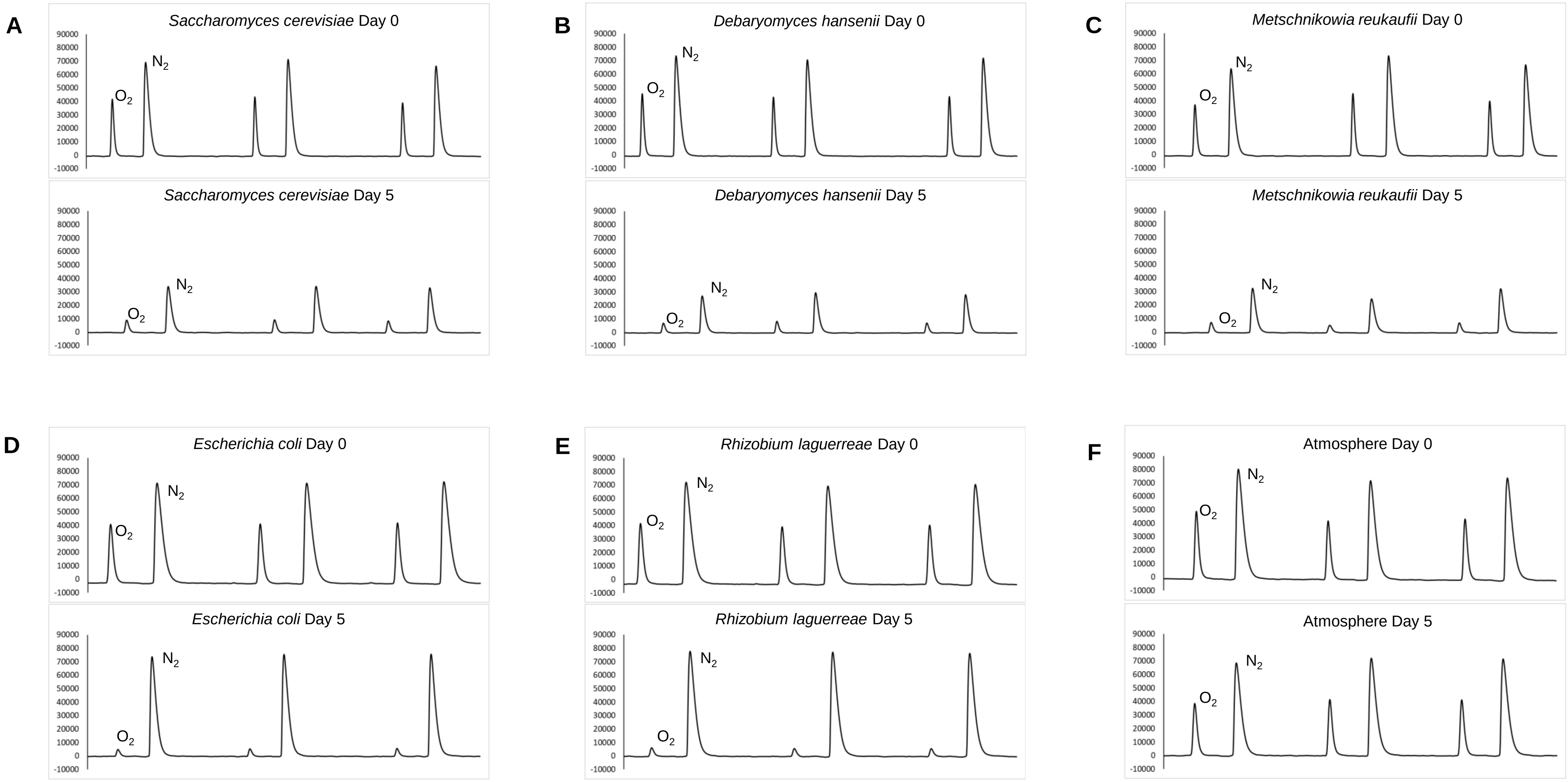


**Suppl. Figure 2**. GC-TCD chromatograms of $N_2$ and $O_2$ in closed liquid cultures in YPD medium at Day 0 and Day 5 after the inoculation: **A.** of Saccharomyces cerevisiae; **B.** of *Debaryomyces hansenii*; **C.** of *Metschnikowia reukaufii*; **D.** negative control *E. coli*; **E.** negative control *Rhizobium laguerreae*; **F.** negative control Atmosphere. Three independent replicates are presented in each panel. Y axis: Intensity.

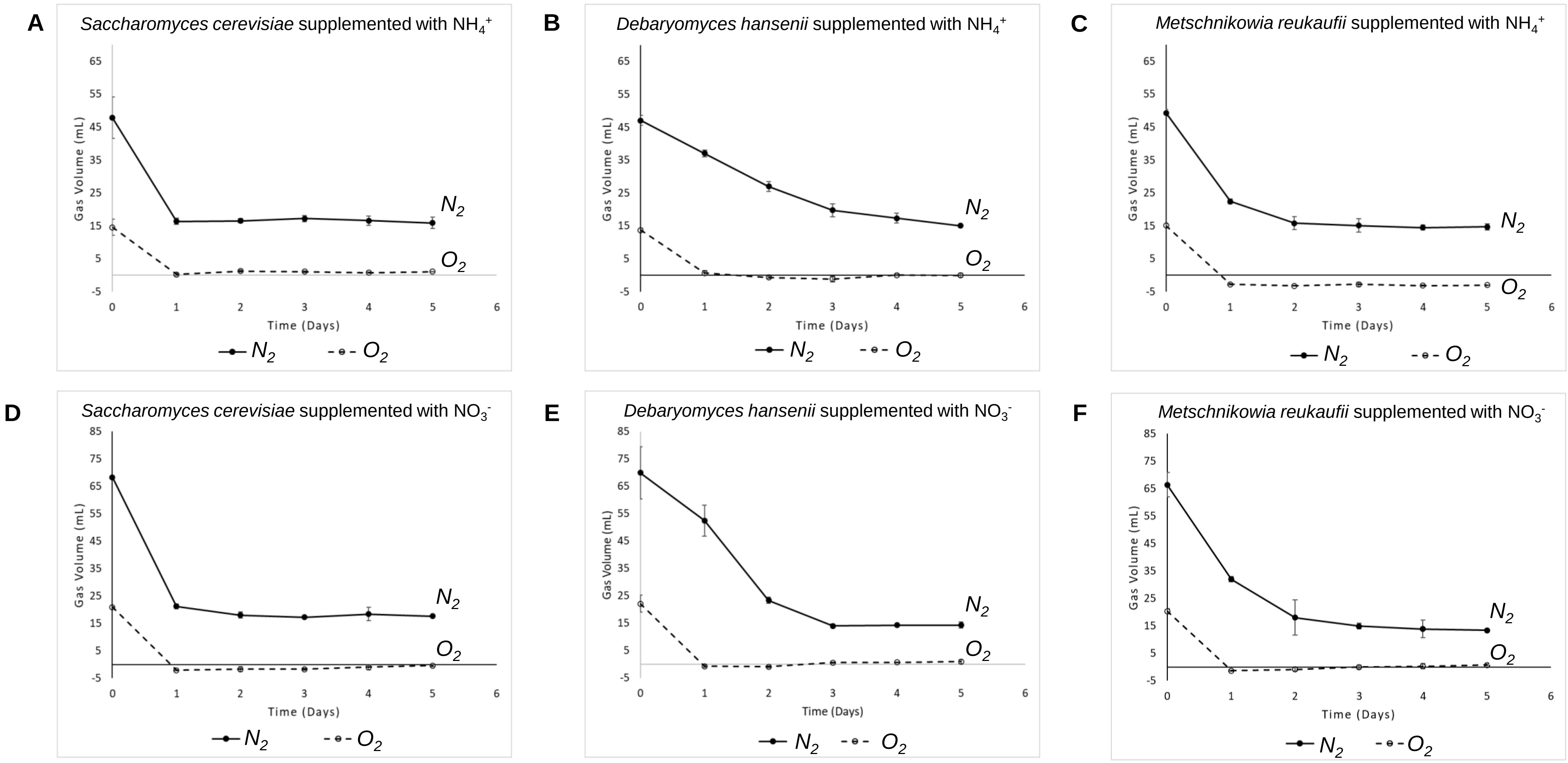


**Suppl. Figure 3**. GC-TCD measurements of $N_2$-assimilation (solid line) and $O_2$ (dashed line) in yeast closed liquid cultures in YPD medium supplemented with $NH_4^+$ or $NO_3^-$: **A.** *Saccharomyces cerevisiae* supplemented with $NH_4^+$; **B.** *Debaryomyces hansenii* supplemented with $NH_4^+$; **C.** *Metschnikowia reukaufii* supplemented with $NH_4^+$; **D.** *Saccharomyces cerevisiae* supplemented with $NO_3^-$; **E.** *Debaryomyces hansenii* supplemented with $NO_3^-$; **F.** *Metschnikowia reukaufii* supplemented with $NO_3^-$.

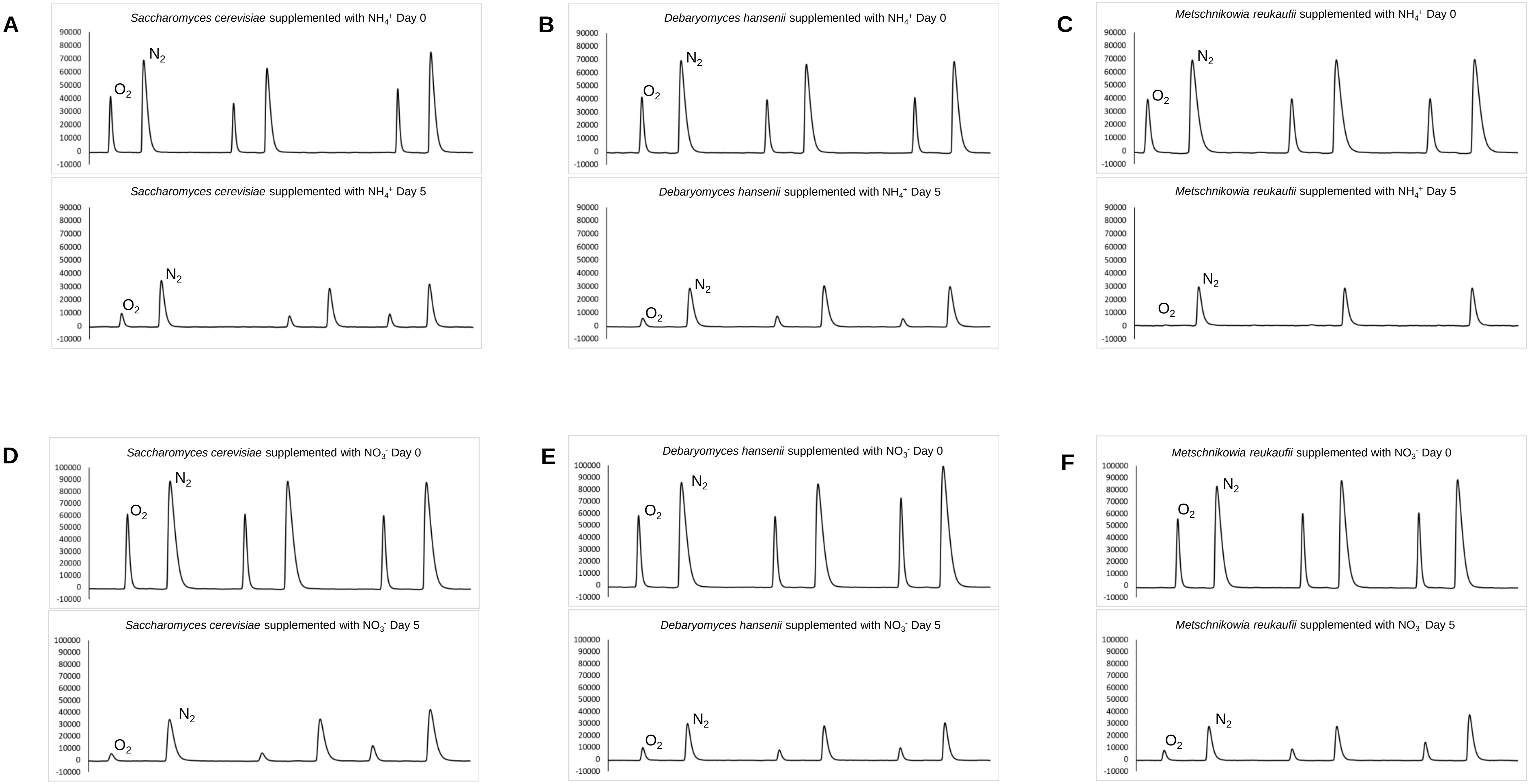


**Suppl. Figure 4**. GC-TCD chromatograms of $N_2$ and $O_2$ in yeast closed liquid cultures in YPD medium supplemented with $NH_4^+$ or $NO_3^-$ at Day 0 and Day 5 after the inoculation: **A.** *Saccharomyces cerevisiae* supplemented with $NH_4^+$; **B.** *Debaryomyces hansenii* supplemented with $NH_4^+$; **C.** *Metschnikowia reukaufii* supplemented with $NH_4^+$; **D.** *Saccharomyces cerevisiae* supplemented with $NO_3^-$; **E.** *Debaryomyces hansenii* supplemented with $NO_3^-$; **F.** *Metschnikowia reukaufii* supplemented with $NO_3^-$. Three independent replicates are presented in each panel. Y axis: Intensity.

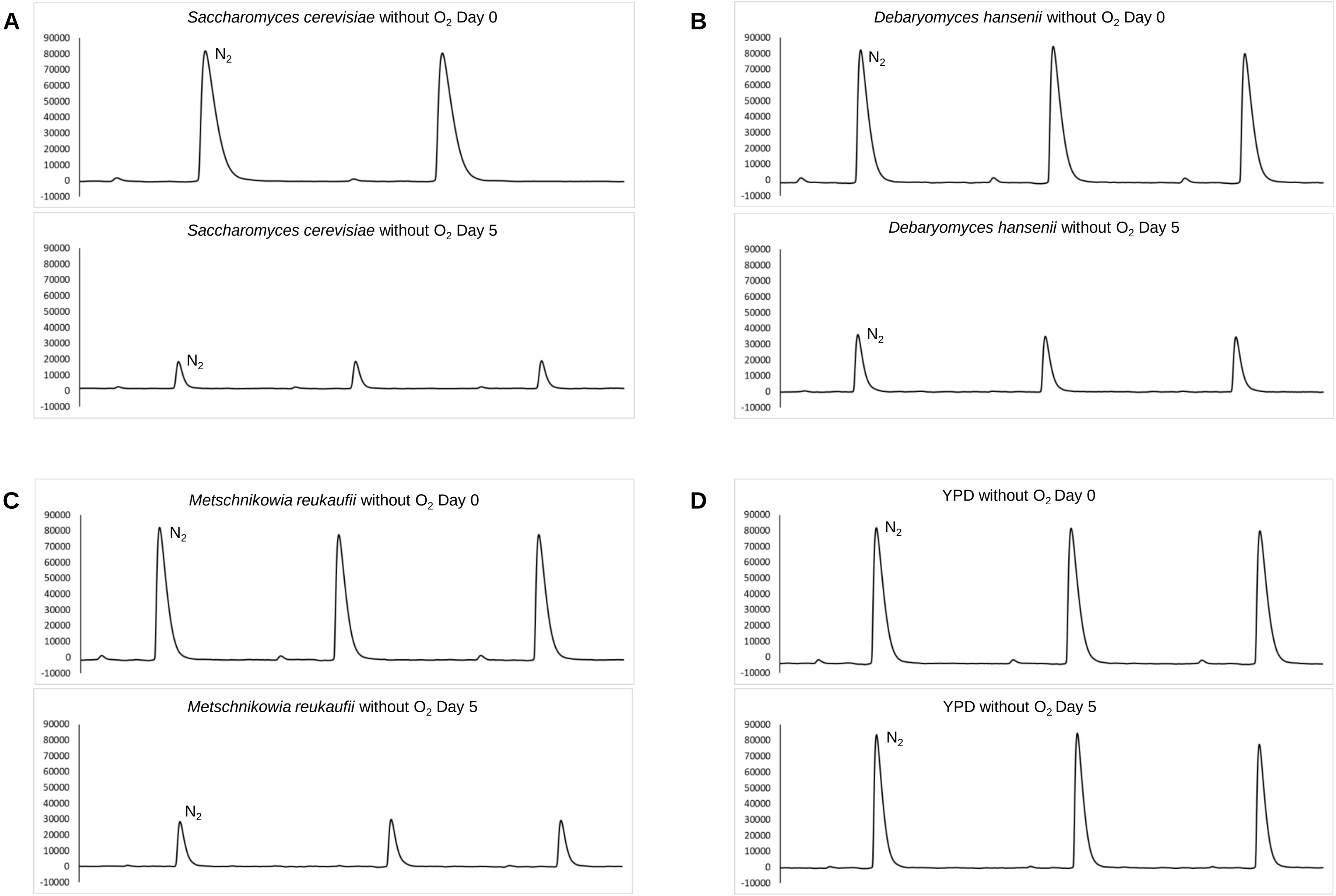


**Suppl. Figure 5**. GC-TCD chromatograms of $N_2$ and $O_2$ in yeast closed liquid cultures in YPD medium without $O_2$ on the headspace at Day 0 and Day 5 after the inoculation: **A.** *Saccharomyces cerevisiae* in YPD medium without $O_2$; **B.** *Debaryomyces hansenii* in YPD medium without $O_2$; **C.** *Metschnikowia reukaufii* in YPD medium without $O_2$ ; **D.** Negative control: YPD medium without $O_2$. Three independent replicates are presented in each panel (two independent replicates are presented for *Saccharomyces cerevisiae* in YPD medium without $O_2$ at Day 0). Y axis: Intensity.

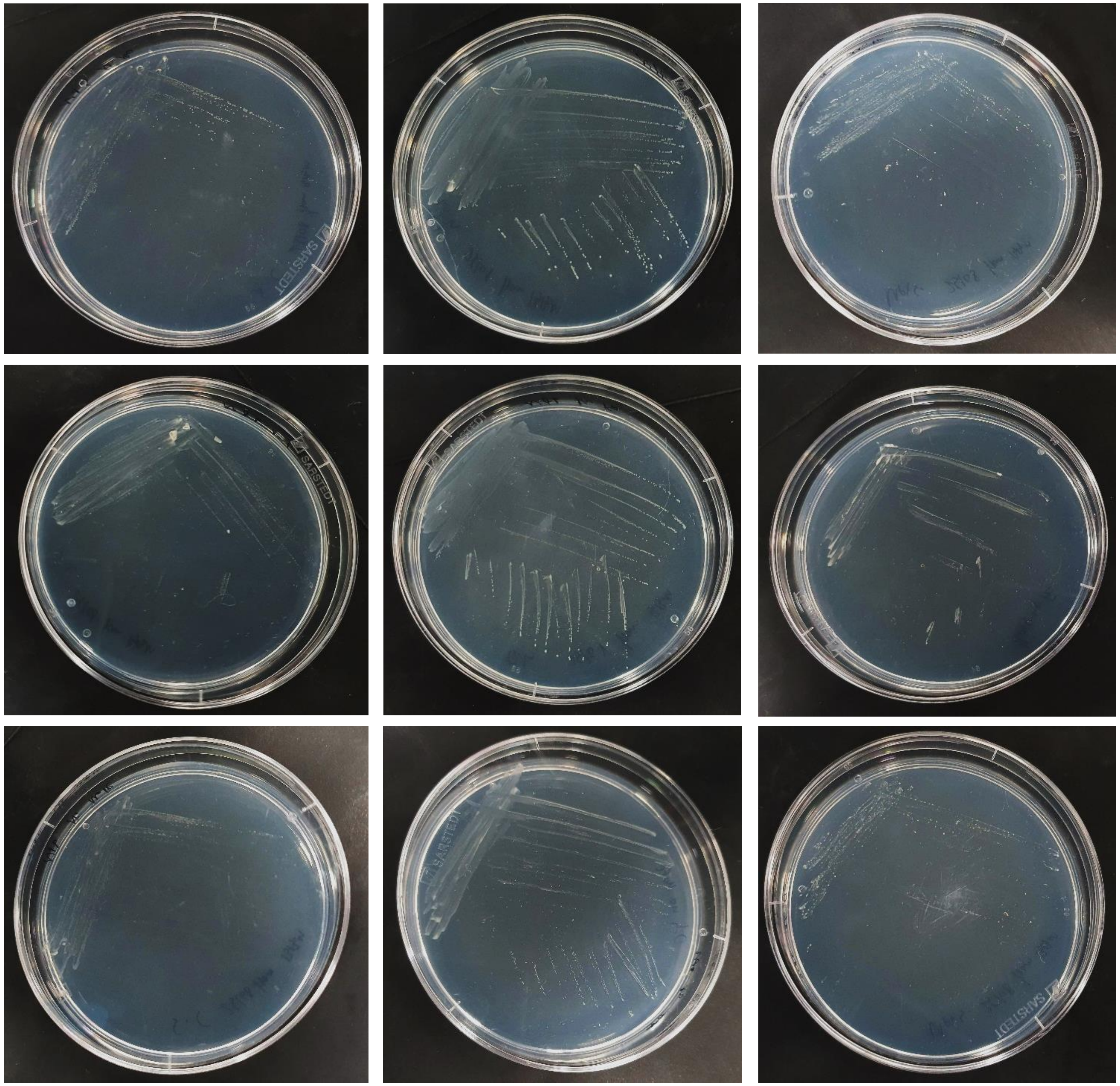


**Suppl. Figure 6**. Growth of all yeast species on nitrogen free NFb medium lacking molybdenum (Mo) sources (NFb w/o Mo), iron (Fe) sources (NFb w/o Fe) and both Mo and Fe sources (NFb w/o Mo and Fe); 4dpi.

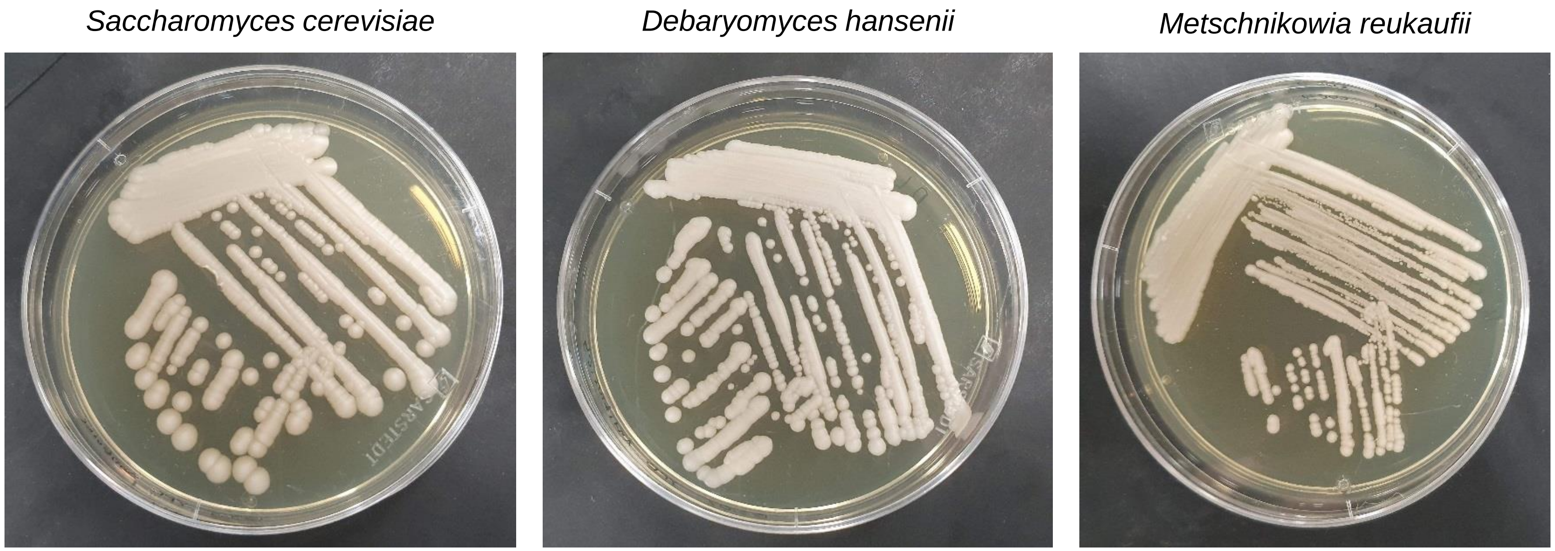


**Suppl. Figure 7**. Growth of all yeast species on reach medium (YPD) supplemented with an antibiotic cocktail; 6dpi.

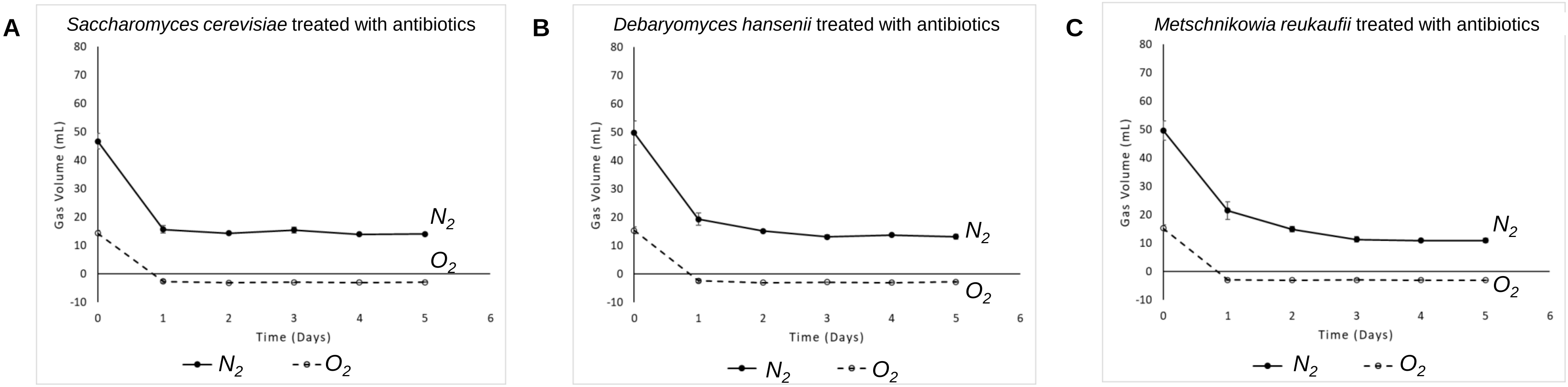


**Suppl. Figure 8**. GC-TCD measurements of $N_2$-assimilation (solid line) and $O_2$ (dashed line) in closed liquid cultures of all yeast species upon treatment with antibiotics: **A.** *Saccharomyces cerevisiae*; **B.** *Debaryomyces hansenii*; **C.** *Metschnikowia reukaufii.*

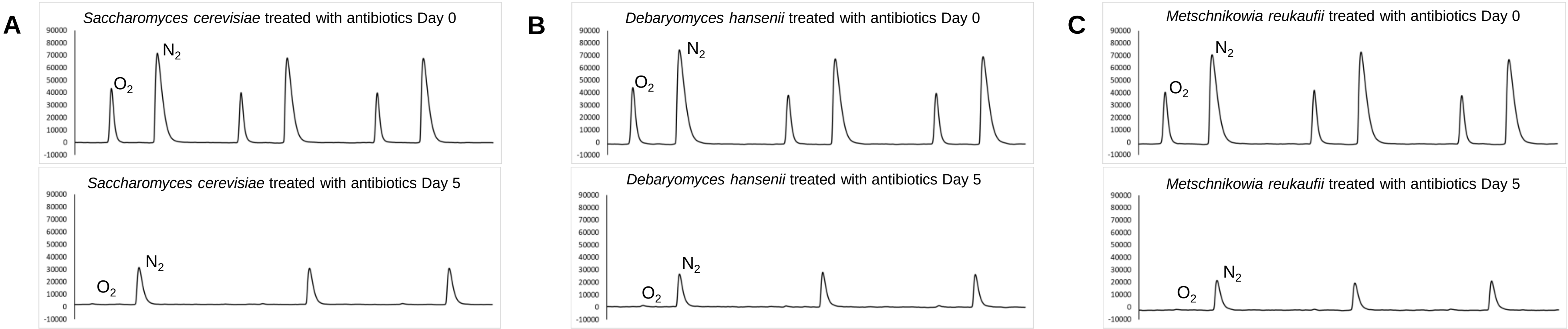


**Suppl. Figure 9**. GC-TCD chromatograms of $N_2$ and $O_2$ in yeast in closed liquid cultures of all yeast species upon treatment with antibiotics at Day 0 and Day 5 after the inoculation: **A.** *Saccharomyces cerevisiae*; **B.** *Debaryomyces hansenii*; **C.** *Metschnikowia reukaufii.* Three independent replicates are presented in each panel. Y axis: Intensity.

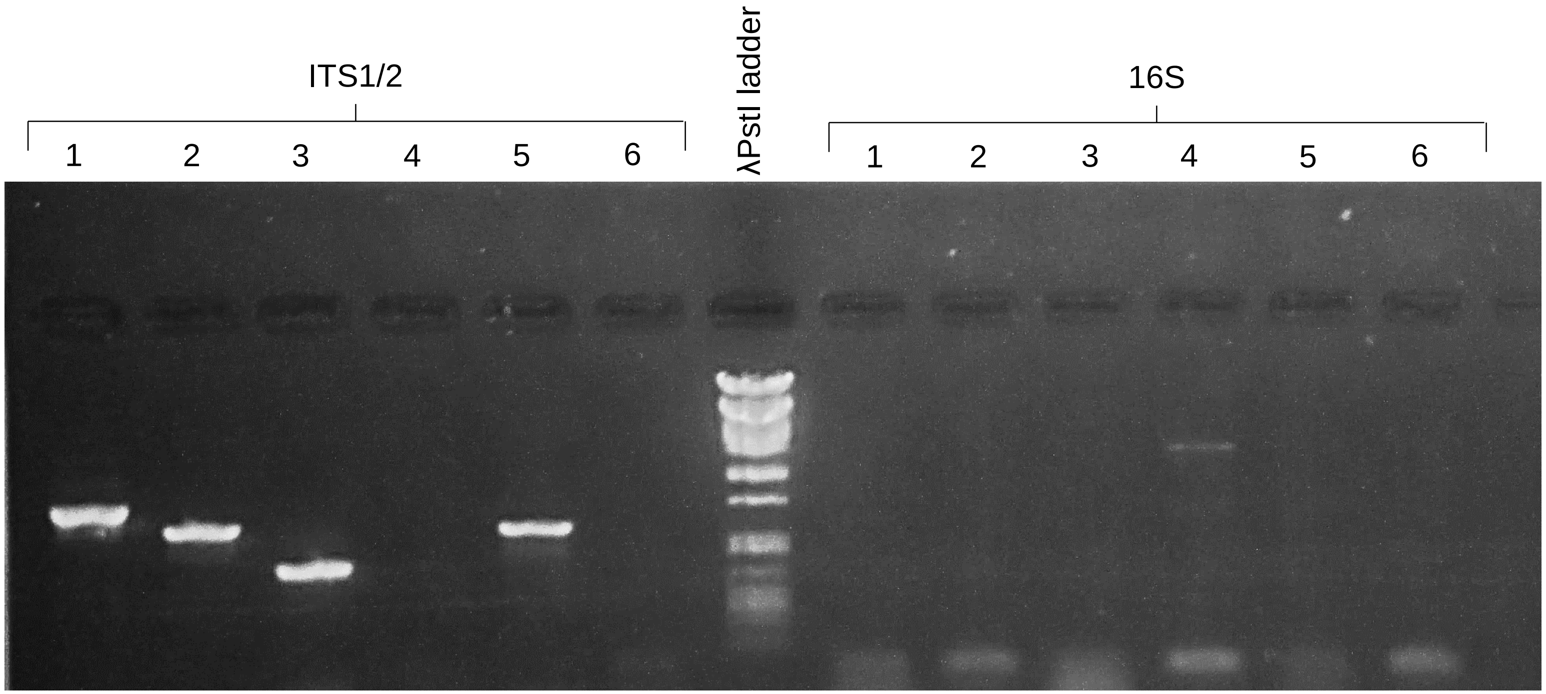


**Suppl. Figure 10**. Electrophoresis of ITS1/2 and 16S diagnostic PCR reactions. ITS1/2: 1. *Saccharomyces cerevisiae*, *2. Debaryomyces hansenii*, 3. *Metschnikowia reukaufii*, 4. *Escherichia coli* (negative control for the ITS1/2 reaction), 5. Fungal DNA (positive control for the ITS1/2 reaction), 6. PCR negative control; 16S: 1. *Saccharomyces cerevisiae*, *2. Debaryomyces hansenii*, 3. *Metschnikowia reukaufii*, 4. *Escherichia coli* (positive control for the 16S reaction), 5. Fungal DNA (negative control for the 16S reaction), 6. PCR negative control.

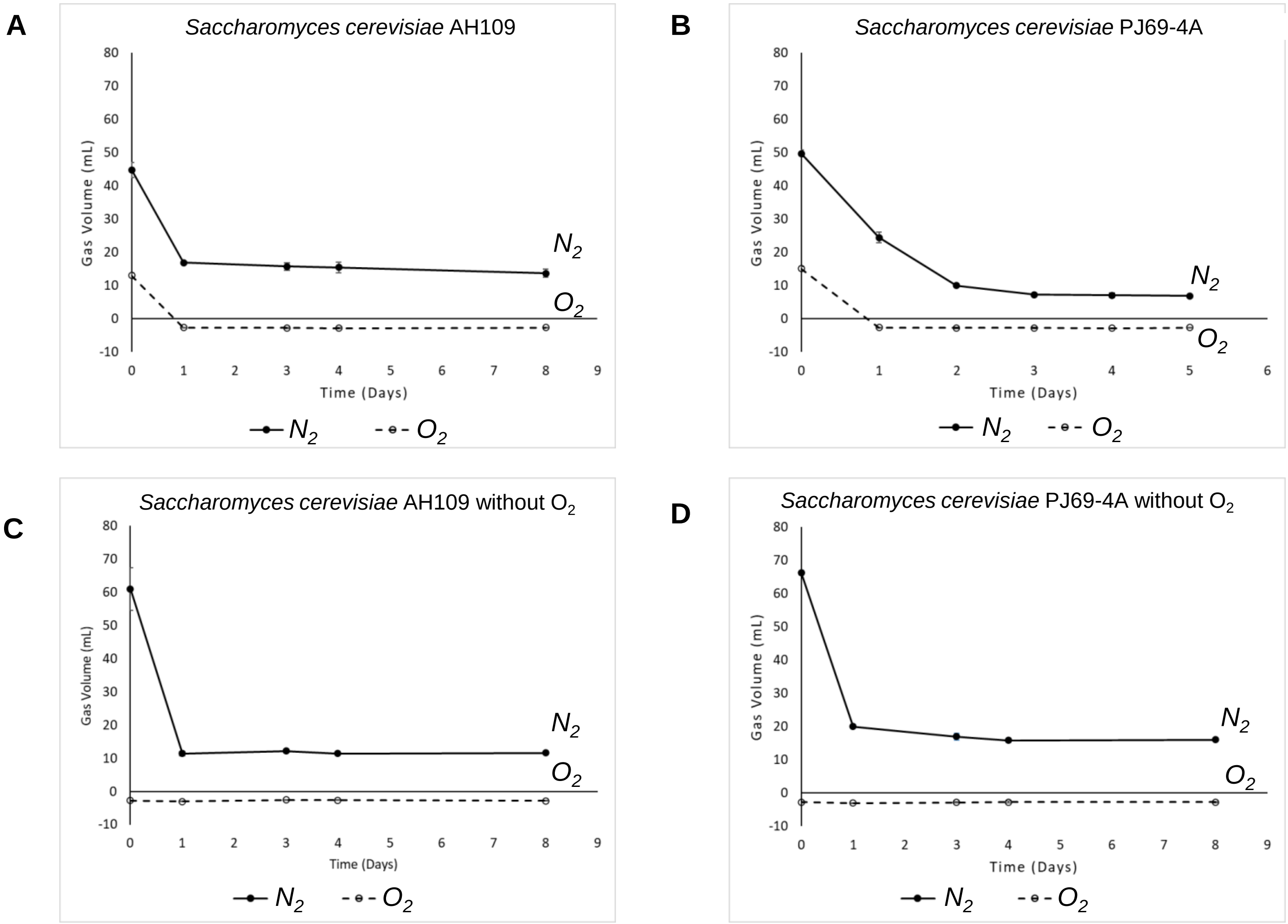


**Suppl. Figure 11**. GC-TCD measurements of $N_2$-assimilation (solid line) and $O_2$ (dashed line) in closed liquid cultures of laboratory yeast strains with and without $O_2$ on the headspace: **A.** *Saccharomyces cerevisiae* AH109; **B.** *Saccharomyces cerevisiae* PJ69-4A; **C.** *Saccharomyces cerevisiae* AH109 without $O_2$; **D.** *Saccharomyces cerevisiae* PJ69-4A without $O_2$.

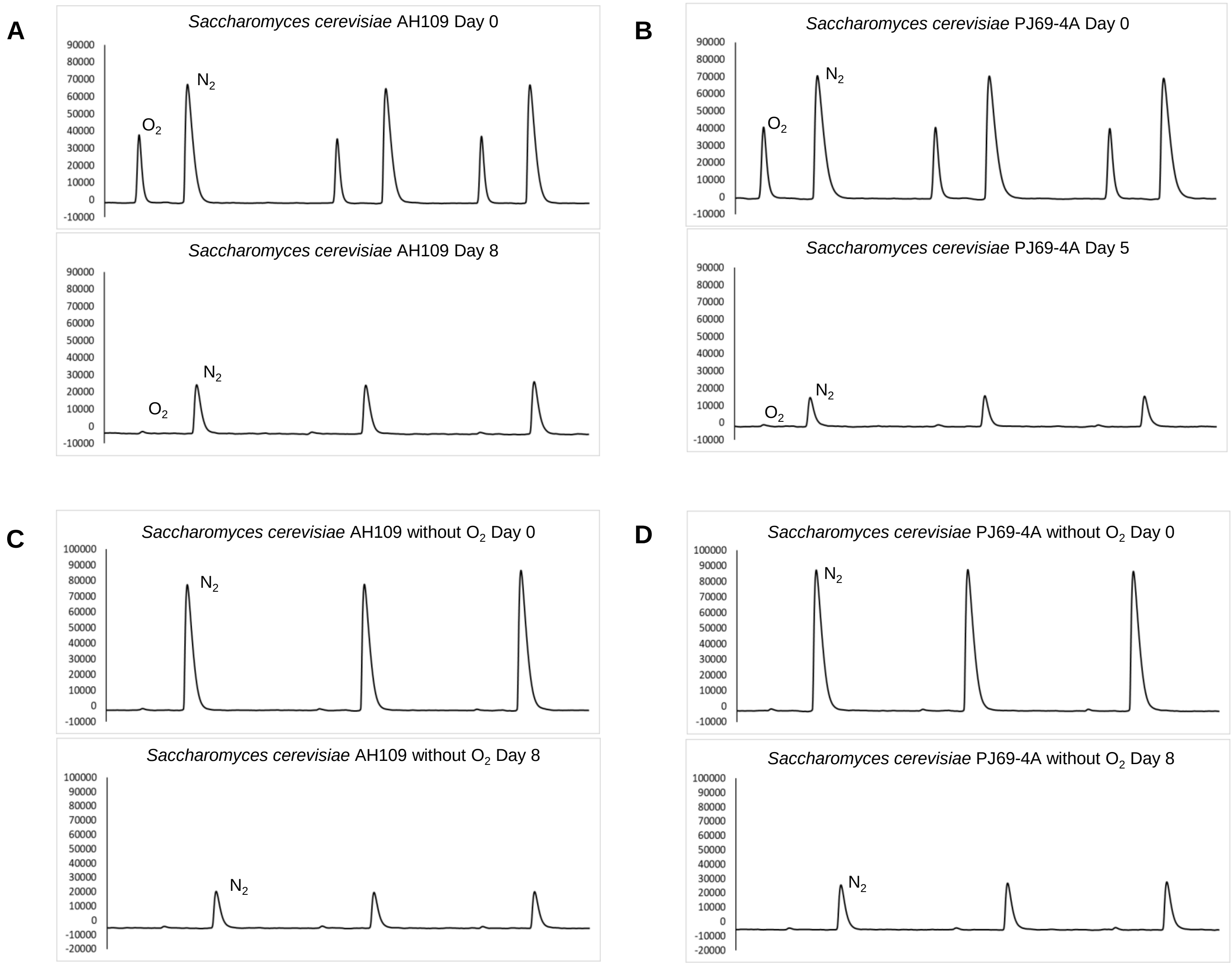


**Suppl. Figure 12**. GC-TCD chromatograms of $N_2$ and $O_2$ in yeast in closed liquid cultures of laboratory yeast strains with and without $O_2$: **A.** *Saccharomyces cerevisiae* AH109 (at Day 0 and Day 8); **B.** *Saccharomyces cerevisiae* PJ69-4A (at Day 0 and Day 5); **C.** *Saccharomyces cerevisiae* AH109 without $O_2$ (at Day 0 and Day 8); **D.** *Saccharomyces cerevisiae* PJ69-4A without $O_2$ (at Day 0 and Day 8).
Three independent replicates are presented in each panel. Y axis: Intensity.
